# Quantifying the effect of forest edge on tropical fauna using explainable ecoacoustics metrics

**DOI:** 10.1101/2025.09.28.679116

**Authors:** Antoine Bouché, Hervé Jourdan, Sylvain Takerkart, Helène de Meringo, Etienne Thoret, Amandine Gasc

**Author notes:** Co-senior authors^b^ (Etienne Thoret, Amandine Gasc).

## Abstract

Tropical forests are biodiversity hotspots but increasingly fragmented by human activities, multiplying forest edges that alter microclimates and species dynamics. While edge effect on plants is well documented, its impact on fauna remains less ex-plored. Ecoacoustics provides a non-invasive and passive approach to monitor faunal communities, yet the complexity of soundscapes and the opacity of machine learning tools often limit ecological interpretability. Here, we show that acoustic diversity in New Caledonian ultramafic forests exhibits a detectable edge effect, with sound-scapes converging towards homogeneity starting 100 m from the forest edge. Using passive acoustic monitoring and 59 ecoacoustic indices, we combined machine learn-ing, Representational Similarity Analysis, and explainable statistical methods from psychophysics. Results reveal a significant correlation between distance to edge and acoustic diversity, particularly pronounced during day-time. Indices such as NDSI, Hf, and SKEWf contributed most to detecting the effect, whereas some other in-dices added noise rather than informative signal. These findings match the botanical threshold described by Blanchard *et al*. (2023) and highlight both the promise and limitations of indice-based ecoacoustics. By integrating explicability methods, we offer a transparent framework to disentangle complex ecological signals, advancing ecoacoustics as a pertinent tool for studying forest fragmentation and guiding biodi-versity conservation.

## I. INTRODUCTION

In a global context of unprecedented environmental change, natural ecosystems are undergoing profound and rapid transformations. The acceleration of global warming and the intensification of human activities through deforestation, urbanization and exploitation of resources are major pressures on biodiversity globally (IPBES, 2019). Forests are reservoirs of biodiversity and are therefore a biological conservation priority (Aerts and Honnay, 2011). Yet, around 10 million hectares of forest disappear annually worldwide (FAO, 2022), changing their structure and affecting their biodiversity. This phenomenon largely contributes to the fragmentation of forests leading to the increase in edge zones, the ecological interfaces between dense forest and open environments. On a global scale, more than 70% of forest areas are located within one kilometer of an edge, and 19% of tropical forests within 100 meters (Haddad *et al*., 2015). Edges are transition zones known to cause significant changes in the microclimate, such as an increase in temperature and a decrease in humidity in the first few meters of the forest (Ewers and Banks-Leite, 2013). These environmental alterations can influence the composition, abundance and behavior of species, thus profoundly modifying the dynamics of biological communities (Haddad *et al*., 2015). This phenomenon is known as the edge effect, defined as a positive or negative variation in the abundance or frequency of a species near the edge, in response to altered environmental conditions and the influence of species or resources from adjacent environments (Driscoll *et al*., 2013). In Amazonian forests, the edge effect on vegetation has already been studied and has been shown to induce profound and lasting changes in forest ecosystems, affecting plant community structure, tree demographic dynamics, and species distribution (Laurance *et al*., 2011).

An emblematic example of forest fragmentation is New Caledonia. It is recognized as one of the world’s 36 biodiversity hotspots, with an exceptional rate of endemism, reaching over 74% for plants (Morat *et al*., 2012). This biological uniqueness is particularly marked in forests on ultramafic soils, i.e., forests that develop on soils rich in heavy metals. These unique environments are home to remarkable biodiversity but are under increasing pressure from human activity such as mining (Birnbaum *et al*., 2023). New Caledonia holds around 25% of the world’s nickel reserves, and mining accounts for over 90% of the territory’s exports (institut de la statistique et des études économiques de Nouvelle-Caĺedonie, 2023). This activity leads to rapid deforestation, particularly in the south of the island where mining exploitation is intensive (Losfeld *et al*., 2015). In this context, it is crucial to better understand the impact of the edge effect on the diversity of forests on ultramafic soils in New Caledonia, in order to assess the consequences of forest fragmentation on local biodiversity. To meet this goal, as part of the RELIQUES project (fRagmentation des forEts sur substrats uLtramafIQUES de Nouvelle-Caĺedonie) (CNRT, 2022), supported by CIRAD and IRD, a first study was carried out in the south of New Caledonia on the response of floristic diversity to the edge effect (Blanchard *et al*., 2023). By combining data on plant composition and canopy structure, this study highlighted clear variations between the first few meters from the edge and areas beyond 100 meters, suggesting a marked ecological gradient linked to fragmentation. However, edge effect studies in general, and the one in New Caledonia in particular, focused on flora (e.g., (Laurance *et al*., 2011)), and more research considering edge effect on fauna are needed. As part of the project RELIQUES, the work presented in this paper complements the floristic analysis conducted by (Blanchard *et al*., 2023) by focusing on the fauna.

### A. Understanding the impact of the edge effect on the fauna’s dynamic using ecoacoustics

Being able to decipher the mechanisms linking fauna’s dynamics to forest density properties necessitates effective monitoring methods that can be applied on a large scale. The ecoacoustic approach (e.g., (Sueur and Farina, 2015)) offers continuous data over long periods of time on a very fine temporal and spatial scale. The recording and analyzing of the sounds of the environment allows a better understanding of interactions between species, their distribution, and the impact of environmental disturbances on biodiversity, ecological dynamics and the overall functioning of ecosystems (Ross *et al*., 2023). A specific application of this ecoacoustic approach is the global analysis of the soundscape of a natural environment. This soundscape, defined by the link between a landscape and the sounds that compose it ((Krause, 1987); (Pijanowski *et al*., 2011)), is divided into three distinct categories: biophony, which groups together all the sound signals emitted by living organisms, geophony, which corresponds to sounds from natural but non-abiotic sources and anthropophony, which includes sounds of human origin unless speech and vocalizations. These three components provide a complete picture of sounds from an ecosystem, and therefore, to some extent, of the composition and state of that ecosystem. In this study, we analyse the soundscapes of New Caledonian ultramafic forests to assess whether forests edge influences the acoustic diversity of the local fauna.

The analysis of soundscapes can be complex due to the multitude and the interdependency of acoustic parameters such as frequency, amplitude, temporal patterns contained in the recorded data. To address this issue, we relied on acoustic indices, which were developed to summarize and quantify soundscape characteristics. A synthesis of these indices has been proposed by (Sueur *et al*., 2014) and updated by (Buxton *et al*., 2018). These indices aim to quantify different characteristics of the soundscape, such as spectral complexity, ratios between frequency bands, frequency richness, entropy or even the temporal organization of signals (Bradfer-Lawrence *et al*., 2025). The application of these indices has proved useful in various ecological contexts. (Krause and Farina, 2016), for example, highlighted a decrease in acoustic complexity, measured by the Acoustic Complexity Index (ACI) (Pieretti *et al*., 2011) which captures temporal variation in sound intensity within frequency bands that reflects the richness of biological activity, in a Californian forest subjected to climatic disturbances, signaling a decline in avian biodiversity. Similarly, (Gasc *et al*., 2018b) were able to monitor the resilience of a desert ecosystem after a fire, by observing the recovery of animal sound signals using the Bioacoustic Index (BI) (Boelman *et al*., 2007) which estimates biotic activity by quantifying acoustic energy in frequency ranges typically occupied by animal sounds. However, a growing number of studies suggest that acoustic indices, when used in isolation, have limited value for monitoring biodiversity, reporting inconsistent performance and variable effect sizes when correlated with biodiversity measurements (Alcocer *et al*., 2022). The results obtained in a certain geographical area would not be reproducible in another area. They would also be difficult to generalize (Sethi *et al*., 2023) and would suffer from poor ecological interpretability. From a methodological point of view, this has led to increasing interest in approaches that consider multiple indices simultaneously.

### B. Explaining black-box statistical methods in meaningful ecoacoustic indices

Analytical approaches from the fields of multivariate statistics and artificial intelligence can integrate several indices at once and detect complex relationships between acoustic indices and ecosystem properties (Thoret *et al*., 2020). However, these methods are often criticized for their opacity. While they allow to conclude that there is relevant information in the acoustic data, it remains challenging to determine which acoustic features drive their predictions or correlation. To overcome this challenge, our study combine advanced statistical methods with explicability techniques specifically designed to address the “black box” nature of these approaches. Following the framework proposed by (Thoret, 2023; Thoret *et al*., 2024, 2021a), we apply explicability methods that help identify the most influential acoustic indices along the distance-to-edge gradient. By incorporating these innovative approaches, our study contributes not only to the understanding of acoustic responses to the distance to edge but also promotes the development of more transparent and generalizable methods in ecoacoustics, an important need in the field.

**FIG. 1.**
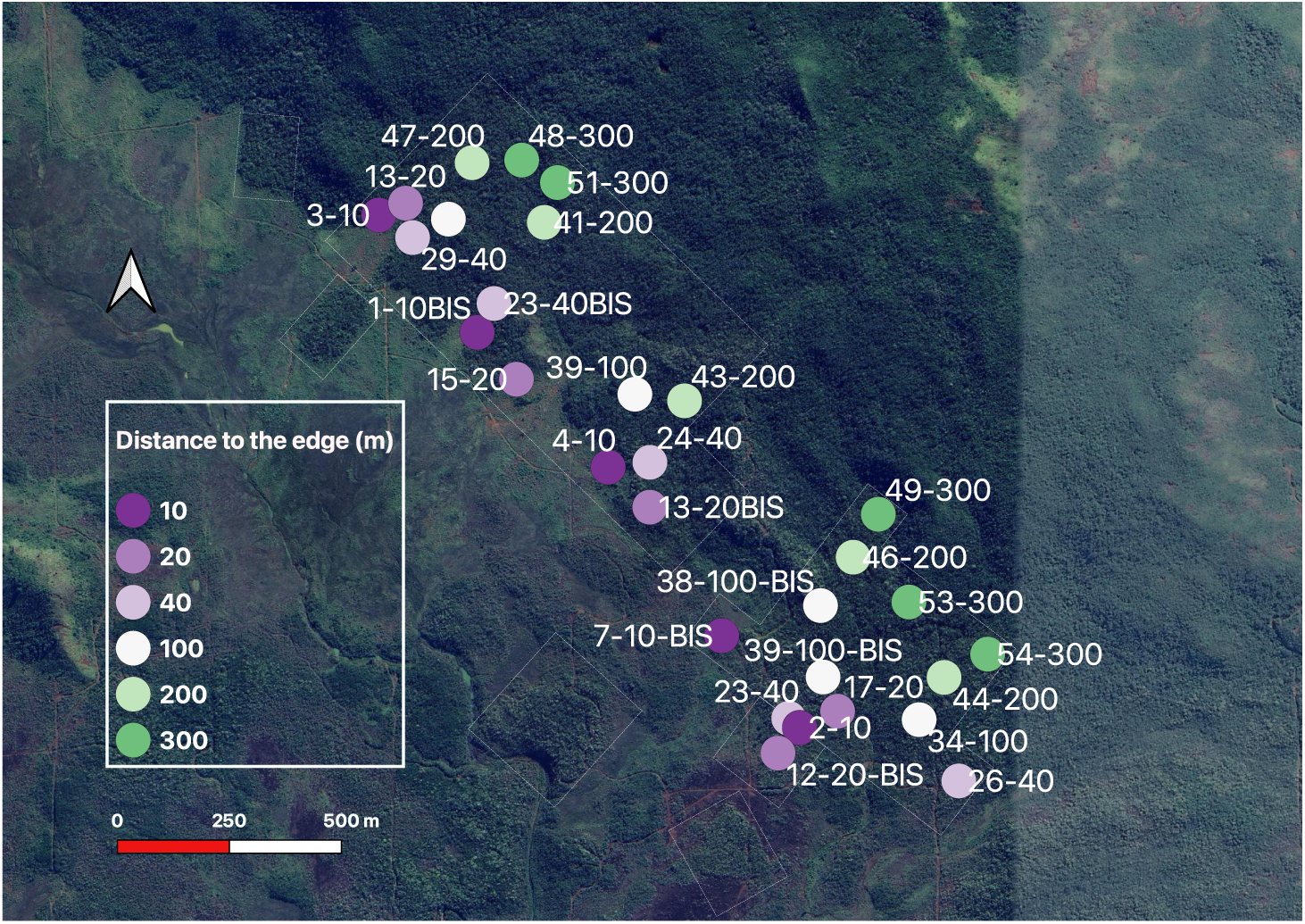
The 30 sites selected for acoustic surveys. 22°13’00“ S - 166°57’00” E and 22°13’56“ S - 166°58’00” E, Kuebini region, southern New Caledonia. Each site is associated to a color point reflecting the distance to the edge and a label corresponding to a unique site name.

## II. MATERIAL AND METHODS

The acoustic dataset was first prepared to ensure that it accurately reflects the biological component of the soundscape and was organized into hourly intervals. Then, we employed a multi-index approach to evaluate the overall impact of edge distance on acoustic diversity. Finally, we used explicability methods to determine which indices contributed most to the observed patterns, revealing which acoustic indices are most important for understanding ecological responses to the edge.

### A. Study sites selection

We selected a large remnant of rain forest at low altitude on ultramafic soils, situated in the southern part of New Caledonia, in the Kuebini Valley region. This kind of low altitude forested area is among the most threatened forest remnants in New Caledonia, as it represents biodiversity relict and conservation refugia, of previously larger forested blocks, that has been fragmented by old past wildfires. This kind of forest is among the most affected by the fragmentation and represent refugia for numerous endemic plants and related fauna, which have all been filtered by harsh conditions from ultramafic (low nutriment, high level of heavy metals in environment). It is situated in Plaine des lacs region where it represents one of the largest patches of free access rain forest. We monitored a total of 30 forested plots (1). For each site, the distance to the forest edge was determined thanks to the analysis of high-resolution satellite images (Blanchard *et al*., 2023). Sites were selected to represent six distances from the forest edge: 10, 20, 40, 100, 200, and 300 meters.

### B. Passive acoustic monitoring

At each site, a SM4 recorder device (Wildlife Acoustics©) was installed on a trunk at 1.50 m from the ground, near the center of the previously established botanical plot (Blanchard *et al*., 2023). The left microphone was systematically oriented towards the north to ensure consistency in the recordings. The device was programmed to record a 1-minute sequence every 10 minutes, to capture variations in the soundscape throughout the day. The recording program was configured as follows: sampling frequency 48000 Hz, gain 16 dB, pre-amplification 26 dB, high-pass filter 220 Hz. All devices were set to UTC+11 (local time) and deployed for at least three weeks, with a minimum target of six consecutive days of favorable weather conditions (no rain or strong wind). Three recording sessions were carried out with ten microphones installed per session in order to have a balanced collection of recordings from the six distances for each session. The first recording session ran from 18 November 2021 to 7 December 2021, the second from 7 December 2021 to 10 February 2022, and the third from 30 June 2022 to 28 July 2022. The three sessions generated a total of 141755 audio files, respectively 17103, 85161 and 39491 files for each session. The variation in the number of files per session is explained by challenges encountered in the field, including weather conditions for collecting the acoustic sensors. Audio files were initially recorded in WAV and then stored in the lossless compression FLAC (Free Lossless Audio Codec) format. This collection is available under the project name RELIQUES from the IMBE sound library (https://ecophony.imbe.fr).

### C. Acoustic indices calculation

For each audio file, we extracted 59 acoustic indices using the *all spectral features()* and *all temporal features()* functions in the Python scikit-maad library (Ulloa *et al*., 2021). These indices provide a fine-grained quantification of the soundscape’s characteristics, based on several signal dimensions: frequency, time, energy, regularity, and spectral diversity. Spectral indices describe how acoustic energy is distributed across the frequency spectrum. For example, the Acoustic Diversity Index (ADI) measures the distribution of energy between frequency bands, while the Acoustic Evenness Index (AEI) reflects the dominance of a limited number of frequencies. Others, such as the Number of Peaks (NBPEAKS), count energy peaks in the spectrum, while spectral entropy measures (e.g., HPairedShannon, HSimpson, Hrenyi) quantify the complexity or predictability of frequency distribution. Temporal indices describe signal variations over time. The Acoustic Complexity Index (ACI), for example, measures rapid changes in intensity, often associated with animal vocalizations. Temporal Entropy (Ht) provides information on signal variability over time, while indices such as Signal-to-Noise Ratio (SNR) indicate overall energy and signal clarity. This multi-index approach forms the basis of our assessment of acoustic variations along the distance-to-edge gradient. All indices are analysed together, without a priori selection, in order to explore these variations exhaustively.

### D. Acoustic disturbance filtering

Acoustic indices, although very useful for analyzing soundscapes, are very sensitive to geophonic or anthropophonic sounds, which could bias the analyses. These elements are likely to modify the spectral structure of the signal and can therefore distort the calculation of acoustic indices. To reduce these identified acoustic biases, acoustic files with microphone-related technical malfunctions were manually discarded. We assumed that no significant anthropophonic sound affected our study area due to its isolation; however, a significant proportion of recordings remained affected by acoustic disturbances of environmental origin, mainly due to geophonic sources such as rain and wind. To discard acoustic recordings containing geophony, we developed an automatic classification model for detecting recordings containing wind or rain.

#### Training dataset

660 audio files were randomly selected from the whole database. These files were manually labeled based on direct listening and visual observation of the spectrogram, to indicate the presence or absence of rain and wind. This step provided a reference training dataset from allowing to develop and evaluate the effectiveness of various filtering methods.

#### Development of the model

We developed a dedicated machine learning classification model, inspired by the work of (Stowell, 2022) on the automatic detection of geophony in acoustic recordings. The labeled dataset (660 files) was randomly divided into a training set (80%) and a test set (20%). Acoustic indices were standardized using *StandardScaler()* to ensure comparability across the variables beforehand. A dimensionality reduction was applied through principal component analysis (PCA), retaining components that explained 99% of the total variance. A Support Vector Classifier (SVC) with a nonlinear kernel, a Radial Basis Function (RBF), was trained. RBF’s hyperparameters, the regularization parameter C, and the kernel parameter gamma, were optimized to maximize the generalization of the SVM using *GridSearchCV* in a five-fold stratified cross-validation scheme (*Stratified-KFold*). Intuitively, these two parameters control the influence of individual training points in the radial basis function (RBF) kernel. The couple of values gamma and C leading to the highest generalization on the held out set was finally retained.

#### Testing

To obtain an estimate of the model’s performance and its ability to generalize to new data, the whole process (data splitting, training, cross-validation, and testing) was repeated 100 times with different random partitions between the training and test sets. The average balanced accuracy over the 100 repetitions is used as the final indicator of model performance. Balanced accuracy corresponds to the average between sensitivity (the rate of correct detection of acoustic nuisance) and specificity (the rate of correct detection of nuisance-free recordings). The classifier achieved a balanced accuracy of 94% on the training data and 85% on the test sample. This corresponds to a specificity of 90% and a sensitivity of 80%. This enabled us to effectively filter out recordings with acoustic nuisance from our entire database (141755 files), allowing us to remove 60493 files containing acoustic nuisance and thus constituting an essential preliminary step in ensuring the reliability of subsequent analyses of acoustic diversity as a function of distance from the edge.

### E. Hourly clustering

After filtering out recordings with acoustic disturbances, we investigated the hourly structuring of soundscapes. It has been established that the acoustic profiles of an ecosystem vary throughout the day (Francomano *et al*., 2020), particularly in relation to the activity rhythms of species in New Caledonian forests on ultramafic soils (Gasc *et al*., 2013). Analyzing these temporal patterns is essential to identify periods that concentrate the most informative acoustic signals, and to ensure that potential edge effect is not masked by daily fluctuations in biological activity. To identify possible hourly groupings, we first calculated the average of each acoustic index per hour by aggregating all available recordings. An unsupervised classification was then performed using the K-means algorithm, preceded by data normalization to ensure fair weighting of indices.

Clustering analysis revealed two distinct groups. The first cluster corresponds to a diurnal (“Day”) period, from 5 a.m. to 6 p.m., closely aligned with sunrise and sunset in the study area, while the second cluster represents a nocturnal (“Night”) period, from 6 p.m. to 5 a.m. (2). This partition highlights the strong daily structuring of the soundscapes and was therefore used as a temporal framework for subsequent analyses.

**FIG. 2.**
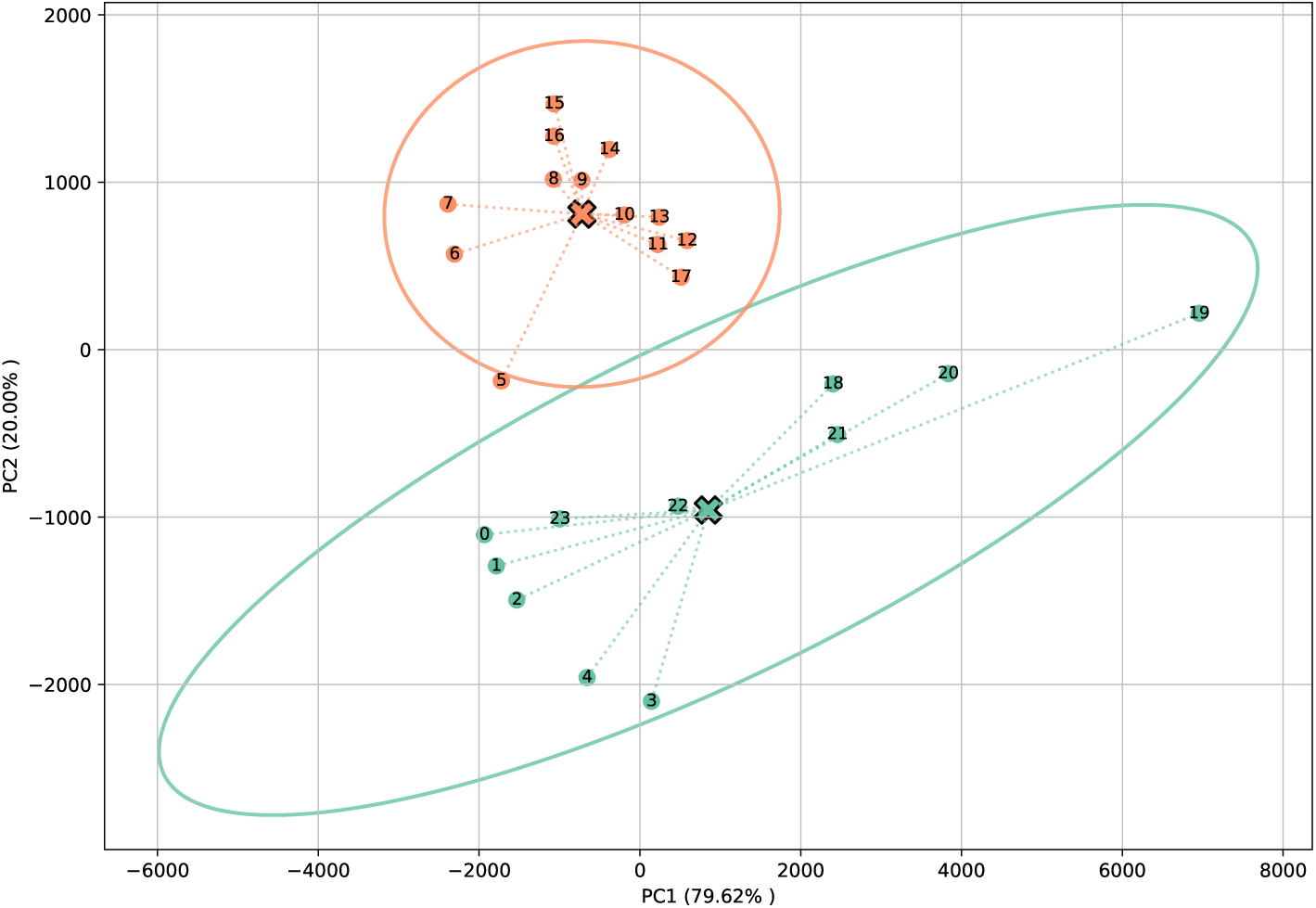
PCA on K-means clustering results. In blue, the “night-time” cluster, In orange, the “day-time” cluster. For each cluster, the center of gravity is represented by a cross and the 95th percentile calculation by an ellipse.

### F. Representational Similarity Analysis

Initially designed to compare neuronal activation patterns, the Representational Similarity Analysis (RSA) introduced by (Kriegeskorte *et al*., 2008) allows analysis of complex structures in multidimensional data sets such as distances matrices. Here we apply this method to evaluate the detection of an edge effect by analyzing the correlations between three dissimilarity matrices designed to quantify different aspects of variation between recordings.

For each pair of recordings, the Euclidean distance was calculated from normalized acoustic indices, producing an acoustic dissimilarity matrix. Two additional matrices were constructed to represent explanatory variables: a “Distance to edge” matrix, indicating the absolute difference in distance to edge for each pair of points, and a binary “Site” matrix, where 0 indicates recordings from the same site and 1 from different sites (3). By correlating the upper triangular part of these matrices with each other, RSA can be used to assess the extent to which acoustic similarity between recordings reflects a particular ecological factor. More specifically, partial correlations were calculated between the acoustic dissimilarity matrix and the “Distance to edge” matrix, using the “Site” matrix as a control variable. This approach isolates the effect of edge distance on acoustic diversity while accounting for the spatial structure of recordings. The analysis can be applied to different time intervals by pre-filtering the relevant matrices, allowing exploration of edge effects over the course of the day.

To build the dataset, six days were randomly selected per site among those with complete recordings covering all precipitation-free time intervals to create a balanced dataset. This choice is based on the work of (Gasc *et al*., 2018a), which indicates that six days are generally enough to characterize the acoustic diversity of a given point in New Caledonian forests of the same area. For each day selected, acoustic indices were averaged over eight three-hour intervals (00-03 h, 03-06 h, …, 21-24 h). This protocol could theoretically generate a total of 1440 records (30 sites × 6 days × 8 intervals). However, to preserve the homogeneity of the dataset, we excluded site 17 due to an insufficient number of valid days, and only four days could be retained for sites 2, 7bis and 29. This gave us a final total of 1368 records that we structured along four hierarchical dimensions: distance to edge, site, day and time interval.

In addition, we investigated at which distance the edge effect is fading. Practically, we investigated at which distance a significant acoustic variation can be detected based using this acoustic dissimilarity matrix. This complete matrix was divided into sub-matrices, each corresponding to recordings from a single distance from the edge. A Pearson correlation was performed between each pair of sub-matrices to assess the similarity of structure between the sites of different distances.

### G. Unveiling relevant ecoacoustic indices of RSA with the “bubbles” method

The RSA method used here is effective in detecting an edge effect based on acoustic characteristics, in this case, ecoacoustic indices. However, it can be described as opaque, as we do not know precisely which indices are decisive or to what extent each contributes to the detection of this effect. It therefore remains difficult to understand exactly which information embedded in the ecoacoustic indices is the most relevant in the sense of RSA correlations.

To improve interpretability of the results, certain methods from the cognitive sciences and psychophysics have recently been adapted to the analysis of machine learning models such as classifiers (Thoret et al., 2021, 2023, 2024) and can be applied to RSA (Giamundo *et al*., 2025). This is the case of the “bubbles” method developed by (Gosselin and Schyns, 2001) in the field of visual perception, to identify the elements of a visual stimulus used by an observer to make a decision, for instance recognizing the gender of someone from their face. The principle consists of randomly masking the stimulus, leaving only small areas called “bubbles” visible so that only certain portions of the information are accessible on each trial. The principle consists of randomly masking the stimulus, leaving only small areas called “bubbles” visible so that only certain portions of information are accessible on each trial. Repeating this procedure makes it possible to estimate which features are most often associated with correct decisions. Applied to ecoacoustics, each feature corresponds to an acoustic index, and its contribution to RSA correlations can be evaluated by perturbing the index vector and testing its effect on correlation scores.

**FIG. 3.**
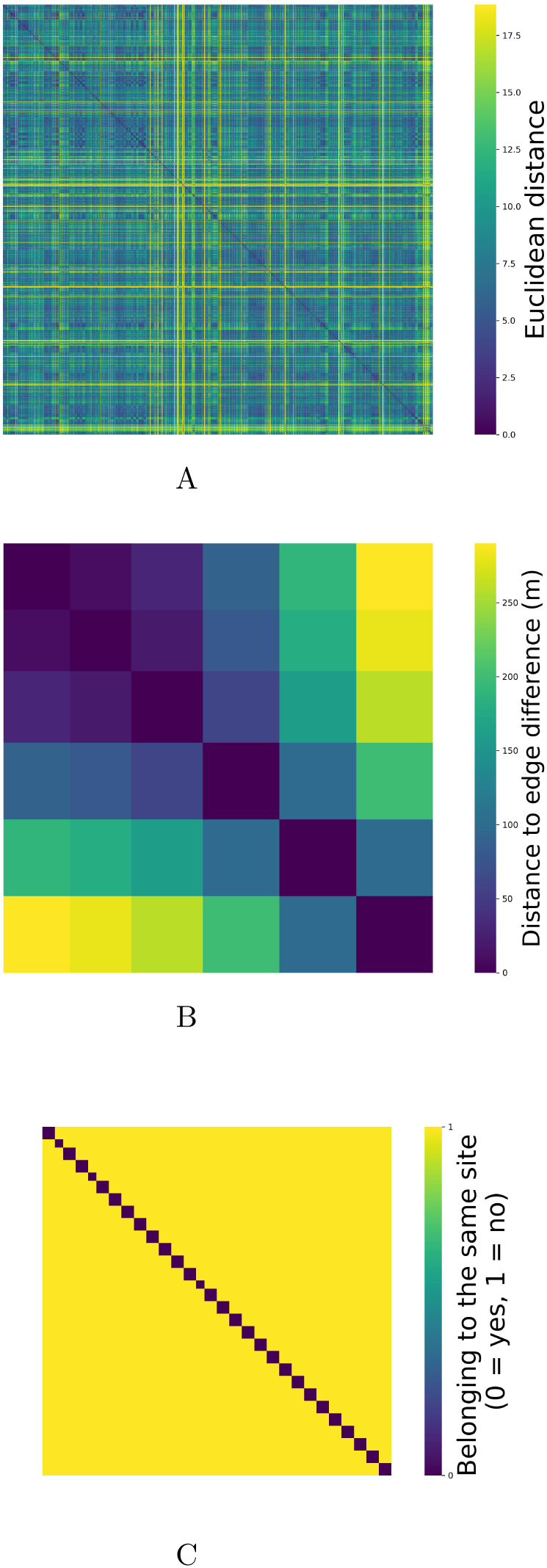
Dissimilarity matrices used in the RSA. (A) “Acoustic” matrix, (B) “Distance to edge” matrix, (C) “Site” matrix.

In practice, we introduce multiplicative noise by randomly removing 15 indices at each iteration and recalculating the partial correlation between the acoustic dissimilarity matrix and the “Distance to edge” matrix, controlling for “Site”. After 10000 iterations, the resulting distribution of correlation coefficients identifies which indices are most decisive in detecting the edge effect, while accounting for redundancy and collinearity common in ecoacoustic indices.

## III. RESULTS

We first analysed the structuring of acoustic diversity with RSA to characterize patterns of homogeneity in deeper forest areas and to evaluate the existence of an edge effect. We then assessed the contrasted contributions of ecoacoustic indices to this effect, identifying the most informative indices and comparing their influence between daytime and nighttime soundscapes.

### A. Influence of forest edge on acoustic diversity

#### 1. Homogeneity of acoustic structures in deeper forest areas

The results suggest a stronger similarity between the acoustic diversity structures at distances of 100 m, 200 m, and 300 m, indicating a certain homogeneity of the soundscapes in these deeper areas of the forest. A complementary analysis was carried out by separating the data according to daytime (05:00-18:00) and night-time (18:00-05:00) periods and showed that the average relationship for distances of 100, 200, and 300 m was 0.063 during the day, compared with 0.033 at night (4).

#### 2. Significant edge effect on acoustic diversity

Results show a significant correlation between the acoustic dissimilarity matrix and the “Distance to edge” matrix: r = 0.0228 (CI95% [0.0206; 0.0247], df = 933658), which is still higher than would be expected under the null hypothesis. This correlation reaches r = 0.0267 (CI95% [0.0220, 0.0306], df = 269742) for the day-time data, while r = 0.0201 (CI95% [0.0157, 0.0242], df = 193750) for the night-time data.

Although the magnitude of these correlations is low, their statistical significance indicates that distance from the edge exerts an influence on the structure of perceived acoustic diversity. These results suggest the existence of an edge effect, albeit a moderate one. It is also possible that the real impact of this effect is partially masked by other factors, such as local environmental variations, biological dynamics that are independent of the edge, or the presence of certain uninformative acoustic indices that introduce noise into the data.

**FIG. 4.**
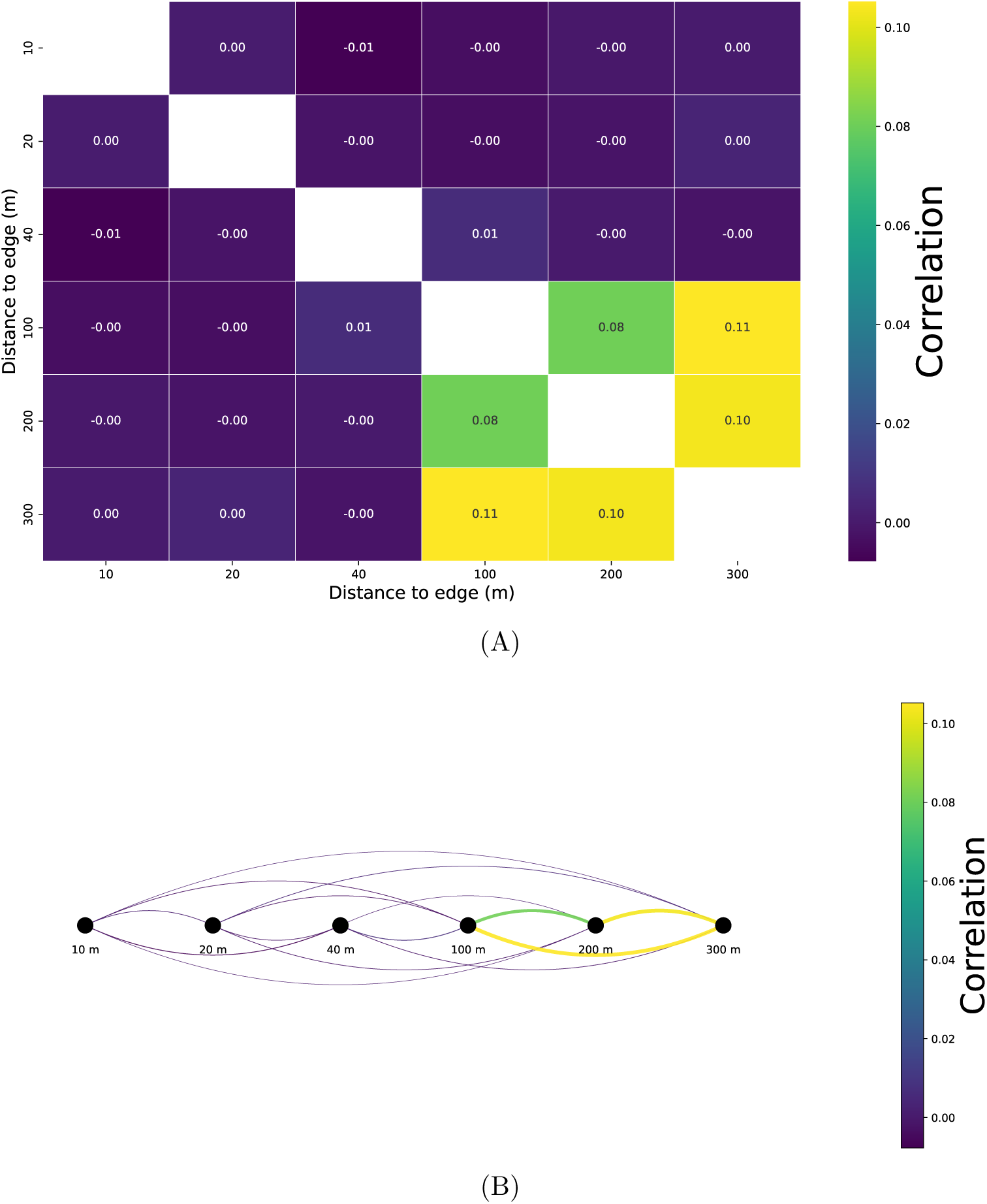
Average correlations between acoustic dissimilarity submatrices according to distance to edge represented by (A) a matrix and (B) a diagram.

Overall, our results indicate that distance from the forest edge affects acoustic diversity. Partial correlation analysis of the three matrices reveals a significant relationship, stronger during daytime, while analysis of only the acoustic dissimilarity matrix shows that sound-scape structures become more homogeneous beyond 100 m, highlighting a detectable edge effect.

### B. Contrasted contributions of ecoacoustic indices on the edge effect detection

Results show that among the 59 indices included in the analysis, 32 contributed positively to the observed correlation while 27 displayed negative weights (5). The most informative indices include NDSI (0.2281), Hf (0.2241), HPairedShannon (0.2136), and SKEWf (0.2005).

By removing these 27 negative indices and recalculating the partial correlation we obtain a coefficient of r = 0.0555 (CI95% [0.0523, 0.0563], df = 933658), a significant improvement compared to the initial score (r=0.0228).

Analysis reveals distinct profiles in the importance of the acoustic indices between the daytime and nighttime periods (5). When only the indices with a positive contribution are retained for each time interval, the partial correlation reaches a coefficient of r = 0.0448 (CI95% [0.0410, 0.0490], df = 269742) for the day-time, compared with r = 0.0564 CI95% [0.0526, 0.0603], df = 193750) for the night-time period.

## IV. DISCUSSION

This study aimed to assess whether an edge effect on fauna acoustic diversity in forests on ultramafic soils in New Caledonia can be detected. Methods from ecoacoustics were combined with multivariate statistics and explainability techniques. A wide range of ecoacoustic indices was calculated from recordings collected at various distances from the edge. These measurements were first used to filter out undesirable recordings with a machine learning classifier trained to detect rain. The relationship between acoustic diversity and distance from the edge was then quantified and characterized. The contribution of different indices was unveiled using correlations between dissimilarity matrices and an explicability method based on reverse correlation. The main finding of our study was to reveal, in New Caledonia forest context, a significant edge effect for animal communities using a biophony analysis, with a detected acoustic activity threshold at 100 meters from forest edge. From a methodological standpoint, this study exemplifies the interest of combining all indices with an explainable pipeline to identify which indices contributed most to detecting this effect and highlighted differences between daytime and nighttime patterns.

**FIG. 5.**
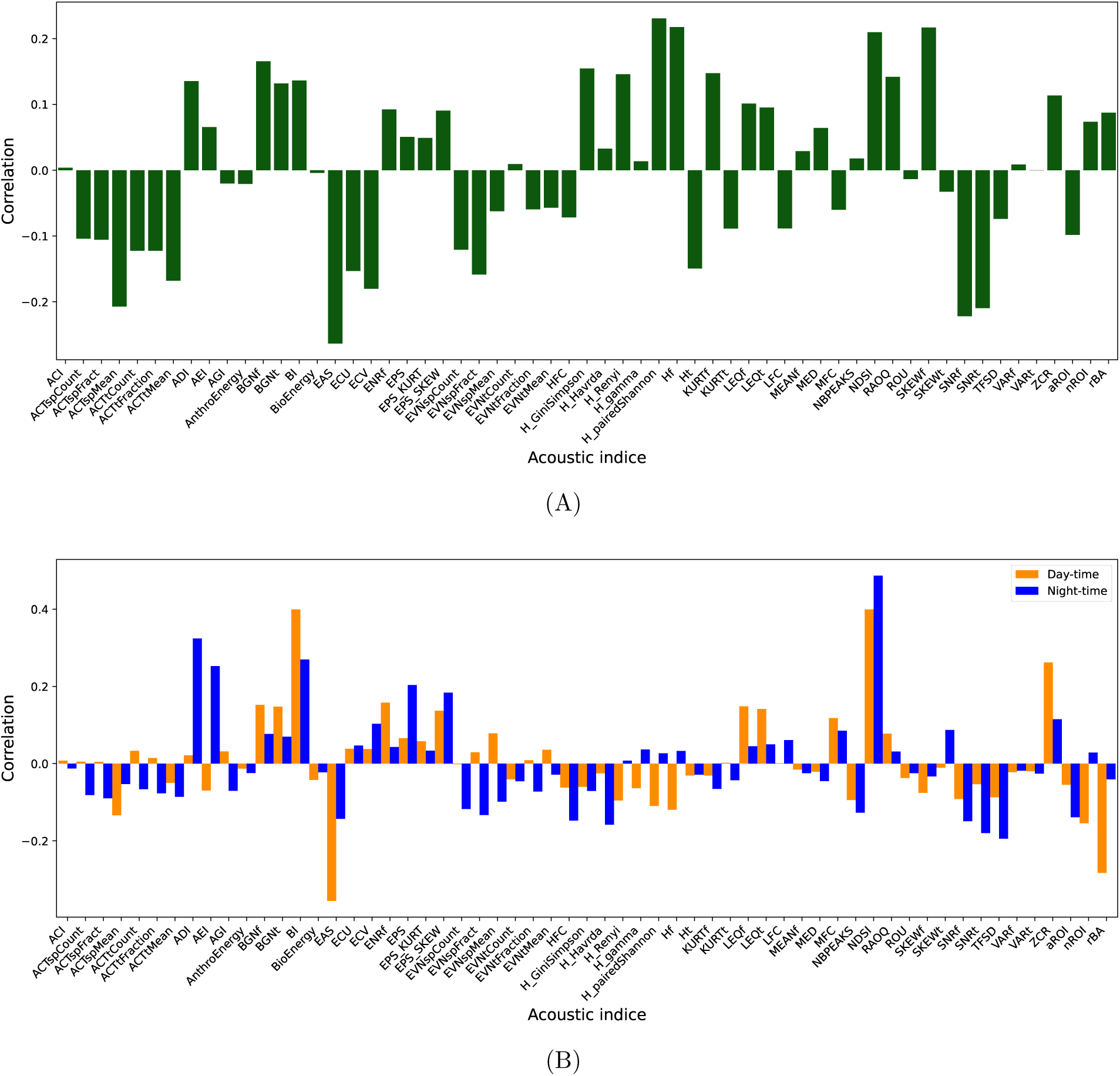
Histogram of the importance of each index in the partial correlation between the acoustic dissimilarity matrix and the “Distance to the edge” matrix, controlling for the site effect. (A) Results obtained by analysing the entire dataset, (B) Results obtained by analysing separately data collected during the “day-time” represented in orange and the “night-time” in blue.

### A. An acoustic edge effect consistent with botanical patterns

The analysis of the acoustic dissimilarity matrices revealed an overall sound structure that was not very similar between the different distances to the edge of the forest, suggesting strong acoustic heterogeneity at the edge of the forest. However, the sites located at 100, 200, and 300 meters show positive correlations between them (r = approximately 0.1), indicating a gradual homogenization of the soundscape beyond 100 meters. These results are consistent with the botanical observations of (Blanchard *et al*., 2023), which also suggest a diminishing edge effect beyond this distance in terms of floristic diversity. They are also in line with the work of (Laurance *et al*., 2011) on tropical forests, where a similar threshold, around 100 meters, was identified for the richness and composition of plant communities. Our result, therefore, supports the existence of a genuine edge effect on faunal activity, perceptible in the overall structure of the soundscape, the intensity of which seems to decrease with distance, potentially reaching a plateau of homogeneity in the deeper areas of the forest. This homogenization can be interpreted as a return to more stable ecological conditions that are less subject to abiotic or human disturbance at the edges.

### B. Other factors contribute to the acoustic diversity

Partial correlation analysis of the dissimilarity matrices was used to isolate the effect of distance from the edge on acoustic dissimilarity, controlling for the site effect. The results show a weak but significant correlation (r = 0.02) when all the acoustic indices are considered. This result suggests that the acoustic structure is influenced by factors other than simple distance from the edge, or that some of the indices considered introduce noise that masks a clearer relationship.

In order to identify the most informative contributions of the indices, the bubble and reverse correlation methods were used. Excluding indices with a negative impact (values below 0) more than doubled the partial correlation compared to the initial value, showing that some indices can add more noise than information in a specific context. This improvement indicates a significant improvement in the analysis, thanks to a data-driven selection of indices.

These results show the importance of selecting indicators beforehand: not all indicators are equally sensitive to the ecological phenomena under study. Their redundancy or sensitivity to noise can alter the quality of the relationships observed, so they must be discarded in order to improve the accuracy of the acoustic analyses. In addition, the information provided by the acoustic indices may vary according to the time interval: the indices used to detect a correlation between acoustic dissimilarity and distance from the edge vary according to the time of day.

Finally, an additional avenue of analysis would be not to exclude the indices associated with a negative impact, but to include them as inverse contributors in the detection of the edge effect. These indices could provide useful ecological information and taking them into account could refine the overall interpretation of the prediction system.

### C. On the interest of combining numerous ecoacoustic indices

The analyses conducted in this study reveal that certain acoustic indices have a high explanatory power with regard to distance from the forest edge. This is particularly the case for the NDSI (Normalized Difference Soundscape Index), Hf (spectral entropy) and SKEWf (spectral asymmetry) indices. Conversely, more commonly used indices such as the ACI (Acoustic Complexity Index) and BI (Bioacoustic Index), which are widely used in ecoacoustic literature, show little contribution here. This divergence highlights an important limitation of approaches based on a small number of indices chosen a priori: they risk obscuring acoustic dimensions that are nevertheless informative in a specific context. The approach adopted here, based on an exhaustive analysis integrating all indices at once and then distilled by explanatory method, allows the selection of truly relevant indices without prior assumptions. However, this ‘data-driven’ approach is not free from bias: some rare indices or those strongly correlated with others may be unfairly excluded. Furthermore, indices considered informative in a given context (e.g. forests on ultramafic soils in New Caledonia) are not necessarily transferable to other ecosystems; this remains to be tested. Thus, combining traditional and explainable approaches appears to be a complementary strategy for refining the ecological relevance of the indices selected.

### D. Explicability and interpretability of models in ecoacoustics

One of the major contributions of this work lies in the use of explainability methods to interpret the results produced by classification methods that are often considered opaque. In the context of ecoacoustics, these models can detect complex relationships between acoustic indices and ecological variables, but without necessarily providing keys to understanding the underlying mechanisms. This is precisely where explainability comes in: by applying the bubble method, it was possible to open the ‘black box’ and determine which acoustic cues are to provide high correlations between matrices. This has not only given us a better understanding of how the soundscape is organized in relation to distance from the forest edge but also revealed which acoustic components convey relevant ecological signals. Explainability is, therefore, a powerful methodological response to the problem of tool opacity, reconnecting the statistical performance of models to a more graspable ecological interpretation.

### E. Limitations

This work has made it possible to characterize the presence of an edge effect on the acoustic soundscape within the Caledonian forests, but several limitations and possible improvements should be highlighted. First, all analyses were based on a single acoustic representation: ecoacoustic indices. Although this innovative approach allows sound information to be condensed into synthetic measurements, it remains reductive compared to the spectro-temporal richness contained in the raw signals. Direct use of representations such as spectrograms, modulation spectra (Chi *et al*., 2005; Elliott and Theunissen, 2009; Thoret *et al*., 2021b) or joint-time frequency scattering (Andén *et al*., 2019) could reveal finer signatures, increase the degree of interpretability by making information associated with distance from the edge more intuitive. Finally, this study focuses solely on the acoustic gradient associated with distance from the edge. A comparative analysis with other gradients, such as botanical gradients (Blanchard *et al*., 2023), should further refine our understanding of the relationship between distance from the edge, the ecological content of the environment, and acoustic variability. This dual perspective, floristic and acoustic, structural and functional, would provide an integrated framework for evaluating the ecological impact of forest fragmentation in New Caledonia.

## V. CONCLUSION

This study explored the effect of distance from the forest edge on acoustic diversity in ultramafic forests in New Caledonia. Thanks to the analysis of ecoacoustic indices geometry with Representational Similarity Analysis, a significant edge effect has been observed. Thanks to an explainable approach, the indices related to this edge effect, NDSI, HPaired-Shannon, Hf and SKEWf have been unveiled. They seem particularly prominent during the daytime, suggesting an increased influence on the daytime acoustic activity of species.

Furthermore, a convergence of acoustic profiles is observed beyond 100 meters from the forest edge, which is consistent with the results obtained by botanical studies conducted in the same area.

Several avenues for further development can be envisaged. First, the use of artificial intelligence models has optimized the processing of a massive dataset, in particular through the automatic filtering of disturbed recordings. Second, the explainability method currently applied to acoustic indices could be extended to the direct analysis of spectrograms. Such an approach would make it possible to identify the precise frequency signatures that characterize different distances from the forest edge. Next, a more detailed hourly analysis, incorporating biological rhythms (such as bird singing periods), could reveal more subtle edge effects that may be masked in aggregated analyses.

Extending the protocol to other ecosystems, with more points per distance, would allow the robustness and transferability of the results to be tested. The goal is to develop automated, reliable and non-invasive monitoring tools to monitor ecological fragmentation via soundscapes.

Finally, this approach opens broader prospects in the field of conservation, particularly in areas that are difficult to access. Acoustic monitoring, a non-intrusive and automatable method, could be particularly relevant in environments such as the marine environment, where traditional ecological monitoring methods are costly or logistically limited.

In this sense, the approach developed here could contribute not only to the ecological management of Caledonian forests, land-use planning and the preservation of endemic biodiversity, but also to a broader transformation of ecological monitoring tools in contexts marked by challenges of resilience to increasing anthropogenic and climatic pressures.

Finally, in the context of Human Auditory Ecology (Lorenzi *et al*., 2023), this study raises the question of the auditory perception of distance to the edge is interesting in itself. This question would be of great interest to be addressed with regard to the knowledge in spatial hearing. Such a context may help decipher how humans might have learned to use acoustic information from natural auditory scene in order to infer information about their environment.

## VII. DATA AVAILABILITY

Data can be accessed upon reasonable request.

## ACKNOWLEDGMENTS

Amandine Gasc and Etienne Thoret co-supervised this work. This work was supported by the project FRagmentation des forEts sur substrats uLtramafIQUES de Nouvelle-Caĺedonie (RELIQUES) directed by Philippe Birnbaum and granted by the CNRT Nickel & son envi-ronment. This work was also supported by grants ANR-16-CONV-0002 (ILCB), ANR-24-CE38-4175 (PSIND), the Excellence Initiative of Aix-University (A*MIDEX), and by CNRS Mission pour l’Interdisciplinarité (MITI) Initiative under the ReScape project, and by the Labex DRIIHM–OHMi Tessekere (French programme “Investissement d’avenir” ANR-11-LABX-0010) for 2024-2026.

We would like to warmly thank Ewan David, Prisca Mahé, Pierre Wallyn, Thomas Cochenille & Valentin Mitran for their valuable help in the field. We also wish to warmly thank for logistic support the team of the Environmental Preservation Department of Prony Resources, especially Stephane Mc Coy. Colleagues from AMAP, IRD, CIRAD Lab & New Caledonian Herbarium, especially Philippe Birnbaum, Gregoire Blanchard & Vanessa Hequet for fruitful discussions on edge effect measurement protocol and the selection of field plots, as well as sharing botanical informations related to these forest plots. Part of this study was funded by the CNRT (National Center for Technological Research) nickel and its environment (www.cnrt.nc) through the RELIQUES project (Grant CSF N° 1PS2017-CNRT /Reliques).

